# The EMT transcription factor Snai1 maintains myocardial wall integrity by repressing intermediate filament gene expression

**DOI:** 10.1101/2020.12.15.422833

**Authors:** Alessandra Gentile, Anabela Bensimon-Brito, Rashmi Priya, Hans-Martin Maischein, Janett Piesker, Stefan Günther, Felix Gunawan, Didier Y.R. Stainier

## Abstract

The zinc finger transcription factor Snai1 is a well-known regulator of epithelial-to-mesenchymal transition (EMT)^1, 2^; it is required for mesoderm ingression in flies^3^ and neural crest delamination in vertebrates^4^. During cardiac development, Snai1-regulated EMT is necessary for myocardial precursor migration and valve formation^5, 6^. However, a role for Snai1 in maturing cardiomyocytes (CMs) has not been reported. Here, using genetic, transcriptomic and chimeric analyses in zebrafish, we find that Snai1b is required for myocardial wall integrity. Global loss of *snai1b* leads to the extrusion of CMs away from the cardiac lumen, a process we show is dependent on cardiac contractility. Examining CM junctions in *snai1b* mutants, we observed that N-cadherin localization was compromised, thereby likely weakening cell-cell adhesion. In addition, extruding CMs exhibit increased actomyosin contractility basally, as revealed by the specific enrichment of canonical markers of actomyosin tension - phosphorylated myosin light chain (active myosin) and the α-catenin epitope α-18. By comparing the transcriptome of wild-type and *snai1b* mutant hearts at early stages of CM extrusion, we found the dysregulation of intermediate filament genes in mutants including the upregulation of *desmin b*. We tested the role of *desmin b* in myocardial wall integrity and found that CM-specific *desmin b* overexpression led to CM extrusion, recapitulating the *snai1b* mutant phenotype. Altogether, these results indicate that Snai1 is a critical regulator of intermediate filament gene expression in CMs, and that it maintains the integrity of the myocardial epithelium during embryogenesis, at least in part by repressing *desmin b* expression.

## Results and Discussion

### The transcription factor Snai1b maintains myocardial wall integrity

As the contractile units of the heart, CMs have to maintain a cohesive tissue-level cytoskeleton to beat synchronously and withstand mechanical force^7, 8^. Using zebrafish as a model to analyze CM cytoskeletal organization at single-cell resolution, we searched for candidate transcription factors that regulate CM cytoskeletal and tissue integrity. Amongst the transcription factors involved in cardiac development, we focused on Snai1, whose orthologues regulate cytoskeletal remodeling and epithelial tissue integrity in *Drosophila* embryos and in mammalian cell culture^9–11^. Snai1 is required for the migration of myocardial precursors during heart tube formation^6^ and for valve formation^5^, but a role in myocardial wall development has not been addressed. Expression pattern analysis revealed the expression of one *snai1* paralogue^12^, *snai1b*, in myocardial and endocardial cells at 50 hours post fertilization (hpf) in zebrafish (data not shown). To analyze *snai1b* function, we generated a promoter-less *snai1b* allele (Figure S1A), which displays almost undetectable levels of *snai1b* mRNA and no transcriptional upregulation of its paralogue^13^ (Figure S1B). Approximately 50% of mutant embryos exhibit cardiac looping defects (Figures S1C–S1D’) as previously reported for *snai1b* morphants^6^. Upon close examination of the *snai1b* mutants with no obvious cardiac looping defects, we observed a new and surprising phenotype leading to a disruption in myocardial wall integrity: CMs were extruding away from the cardiac lumen. We found that both homozygous and heterozygous *snai1b* embryos exhibited a significant increase in the number of extruding CMs compared to their wild-type siblings (Figures 1A–1E). The frequency of this CM extrusion is higher in the atrioventricular canal (AVC) (Figure S1E), where the cells are exposed to strong mechanical tension from the blood flow and looping morphogenesis^14–16^. CM extrusion was observed starting at 48 hpf and continued during larval stages including at 78 (Figures S2A–S2C) and 100 (Figures S2E–S2G) hpf. These results indicate a previously uncharacterized role for Snai1b in maintaining myocardial tissue cohesion.

**Figure 1.**
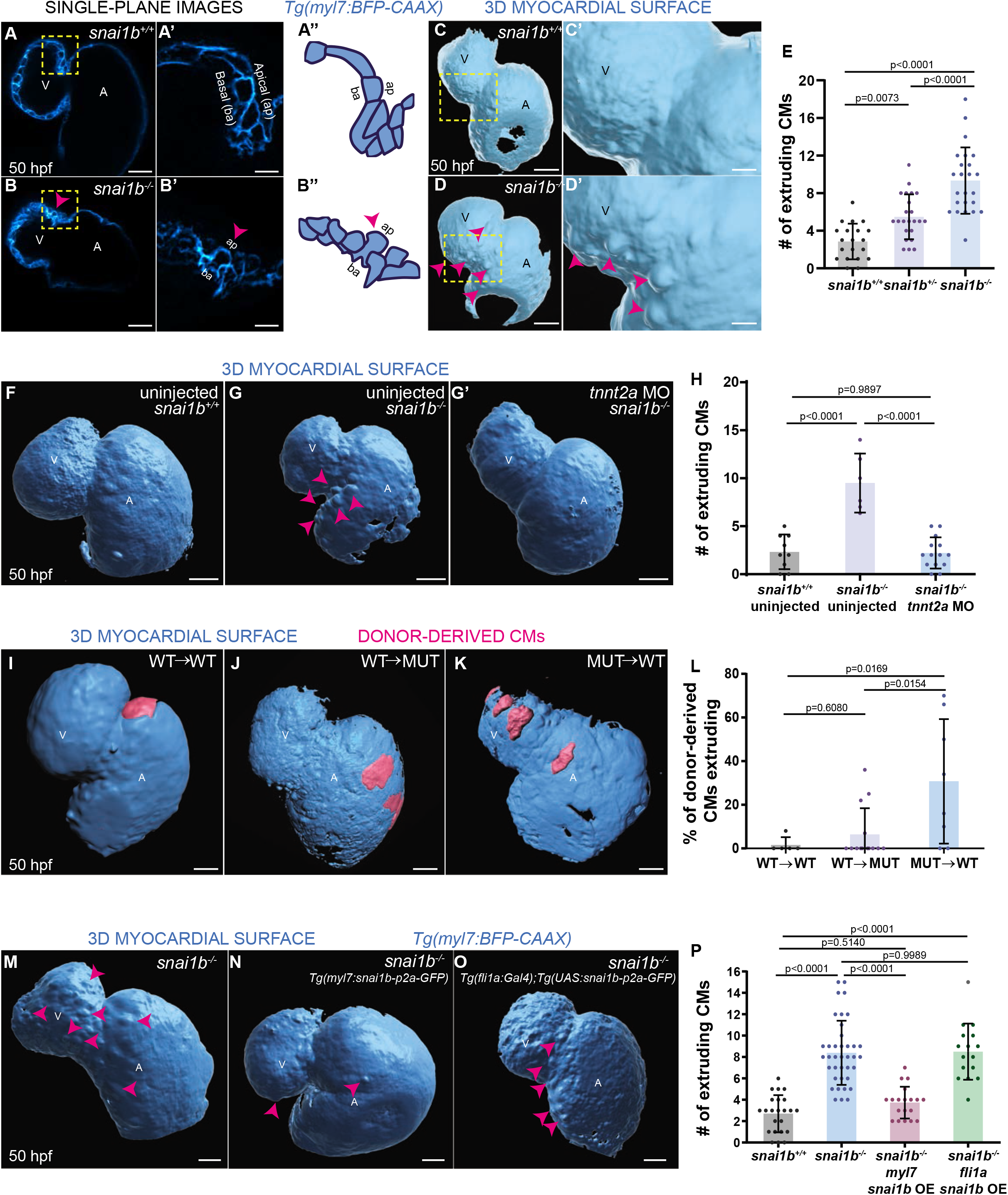
Loss of *snai1b* leads to cardiomyocyte extrusion, disrupting myocardial wall integrity. **A-B”)** Single-plane images of *snai1b*^+/+^ (A) and *snai1b*^-/-^ (B) hearts at 50 hpf. Close-up of boxed areas (A’, B’), and schematic (A”, B”). **C-D’)** 3D surface rendering of the myocardium of *snai1b*^+/+^ (C, C’) and *snai1b*^-/-^ (D, D’) embryos at 50 hpf. CM extrusions are clearly observed in *snai1b*^-/-^ embryos (magenta arrowheads in B, B’, B”, D, D’). **E)** More CMs are extruding in *snai1b* mutants compared with wild-type and heterozygous siblings at 50 hpf (*snai1b*^+/+^, n =20; *snai1b*^+/-^, n =23; *snai1b*^-/-^, n=24). **F-H)** Blocking cardiac contractions with *tnnt2a* MO leads to a reduced number of extruding CMs in *snai1b*^-/-^. F-G’) 3D surface rendering of the myocardium of *snai1b*^+/+^ (F) and *snai1b*^-/-^ (G) uninjected embryos and *snai1b^-/-^ tnnt2a* morphants (G’). **H)** Fewer CMs are extruding (magenta arrowheads in G) in *snai1b*^-/-^ embryos injected with *tnnt2a* MO (n=14) compared with uninjected *snai1b*^-/-^ (n=6) and *snai1b*^+/+^ (n=9) embryos at 50 hpf. **I-L)** 3D surface rendering of the myocardium showing wild-type donor-derived CMs in a wild-type (I) or *snai1b*^-/-^ (J) heart, and *snai1b*^-/-^ donor-derived CMs in a wild-type heart (K). **L)** The percentage of donor-derived CMs that extrude is higher when *snai1b*^-/-^ donor-derived CMs are in wild-type hearts (n=8) than when wild-type donor-derived CMs are in wild-type (n=5) or *snai1b*^-/-^ (n=14) hearts. **M-P)** Overexpression of *snai1b* specifically in CMs partially rescues the CM extrusion phenotype in *snai1b*^-/-^. 3D surface rendering of the myocardium of a *snai1b*^-/-^ embryo (M), and *snai1b*^-/-^ embryos overexpressing *snai1b* under a *myl7* (N) or a *fli1a* (O) promoter. P) Fewer CMs are extruding (magenta arrows in M, N and O) in *snai1b*^-/-^ embryos (n=19) overexpressing *snai1b* in CMs compared with *snai1b*^-/-^ embryos (n=38), and this number is comparable to that in *snai1^+/+^* embryos (n=24). The number of extruding CMs does not change in *snai1b*^-/-^ embryos (n=16) when *snai1b* is overexpressed in endothelial cells. Plot values represent means ± S.D.; p-values determined by one-way ANOVA followed by multiple comparisons with Dunn test (E,H,L,P). Scale bars: 20 μm. V, ventricle; A, atrium; ap, apical; ba, basal; n, number of embryos.

For all further analysis, we decided to focus on the *snai1b* mutants displaying apparently unaffected cardiac looping. We first analyzed whether the extruding CMs in *snai1b* mutants were apoptotic, as dying epithelial cells are frequently removed by extrusion. However, we did not observe a significant difference in the rate of dying cells, as marked by terminal deoxynucleotidyl transferase dUTP nick end labeling (TUNEL), between *snai1b*^+/+^ (Figure S3A) and *snai1b*^-/-^ hearts (Figures S3A’–S3A”), indicating that CM extrusion in *snai1b* mutants is not due to cell death.

We next asked whether the defects in myocardial integrity have an impact on cardiac morphology and function. Interestingly, although we did not observe a significant reduction in CM numbers or ventricular volume at 50 hpf (Figures S3B–S3C), we noted a reduction in ventricular volume at 78 (Figure S2D) and 100 (Figure S2H) hpf. In addition, we observed a decrease in the number of delaminating CMs in *snai1b*^-/-^ larvae at 78 hpf (Figure S2K), resulting in fewer trabecular CMs at 100 hpf (Figures S2J–S2J’, S2L) compared with wild-type siblings (Figures S2I–S2I’). Taken together, these data suggest that the extrusion of CMs caused by the loss of *snai1b* disrupts myocardial wall morphology.

A role for contractility-induced mechanical forces on cardiac tissue stability has recently been reported^17–19^. Hence, we sought to test whether the loss of cardiac contractility would ameliorate the CM extrusion phenotype in *snai1b* mutants, as previously shown for *klf2* mutants^17^. We observed that after injecting a *tnnt2a* morpholino^20^ (Figures 1G–1G’) to prevent cardiac contraction, the number of extruding CMs in *snai1b* mutants at 50 hpf was significantly reduced, and in fact comparable with that in uninjected *snai1b*^+/+^ embryos (Figures 1F–1H). These data indicate that mechanical forces due to cardiac contraction are required for the increased frequency of CM extrusion observed in *snai1b* mutants.

To test whether Snai1b plays a cell-autonomous role in promoting myocardial wall integrity, we generated mosaic hearts by cell transplantation. We observed that donor-derived *snai1b*^+/+^ CMs remained integrated in the *snai1b*^-/-^ myocardial wall (Figure 1J), whereas donor-derived *snai1b*^-/-^ CMs in a *snai1b*^+/+^ heart were significantly more prone to extrude than their wild-type neighbors (Figures 1K–1L). Together, these data indicate that *snai1b* is required in a CM-autonomous manner to maintain myocardial wall integrity. Furthermore, we found that CM-specific, but not endothelial-specific, *snai1b* overexpression rescued the *snai1b*^-/-^ CM extrusion phenotype (Figures 1M–1P and S1F–S1G), further indicating that Snai1b is required in CMs to suppress their extrusion.

### Snai1b limits CM extrusion by regulating the actomyosin machinery

Some CMs delaminate towards the cardiac lumen during the process of trabeculation^21^. During this delamination, they lose their apicobasal polarity and undergo an EMT-like process^22^. We wanted to determine whether the extruding CMs in *snai1b* mutants also lose their apicobasal polarity. Notably, we observed that the polarity marker Podocalyxin remained on the apical side of the extruding CMs in *snai1b* mutants (Figures S3D and S3E’), suggesting that apicobasal polarity is maintained.

Extensive studies in *Drosophila* embryos and in mammalian cell culture have revealed the importance of cell extrusion in limiting tissue overcrowding and eliminating dying cells to maintain tissue homeostasis and/or determine cell fate^11, 23, 24^. Other experiments have shown that a contractile actomyosin ring around the cortex of the cells is necessary for their extrusion^24–26^. Using antibodies against the α-catenin monoclonal epitope α-18^27^, which recognize the open conformation of α-catenin, a mechanosensitive protein, and against phosphorylated/activated myosin light chain (p-myosin), we assessed cellular contractility in extruding CMs in *snai1b*^+/+^ and *snai1b*^-/-^ embryos (Figures 2A–2B” and 2G–2H”). Increased α-18 and p-myosin immunofluorescence intensity was observed at the basal side of extruding CMs in *snai1b*^-/-^ (Figures 2B–2C’ and 2H–2I’) and *snai1b*^+/+^ (Figures 2A–2A’ and 2G–2G’) embryos. As cellular extrusions also involve the rearrangement of cell-cell junctions^28–30^, we assessed the localization of the major CM adhesion molecule, N-Cadherin^31^. We observed an overall reduction in N-cadherin levels in the junctions between CMs in *snai1b* mutants compared with those in wild-types siblings (Figures 2A-B” and 2D–2D’), suggesting that Snai1 regulates N-cadherin localization to stabilize actomyosin tension at the junctions, thereby sustaining adhesion between CMs.

**Figure 2.**
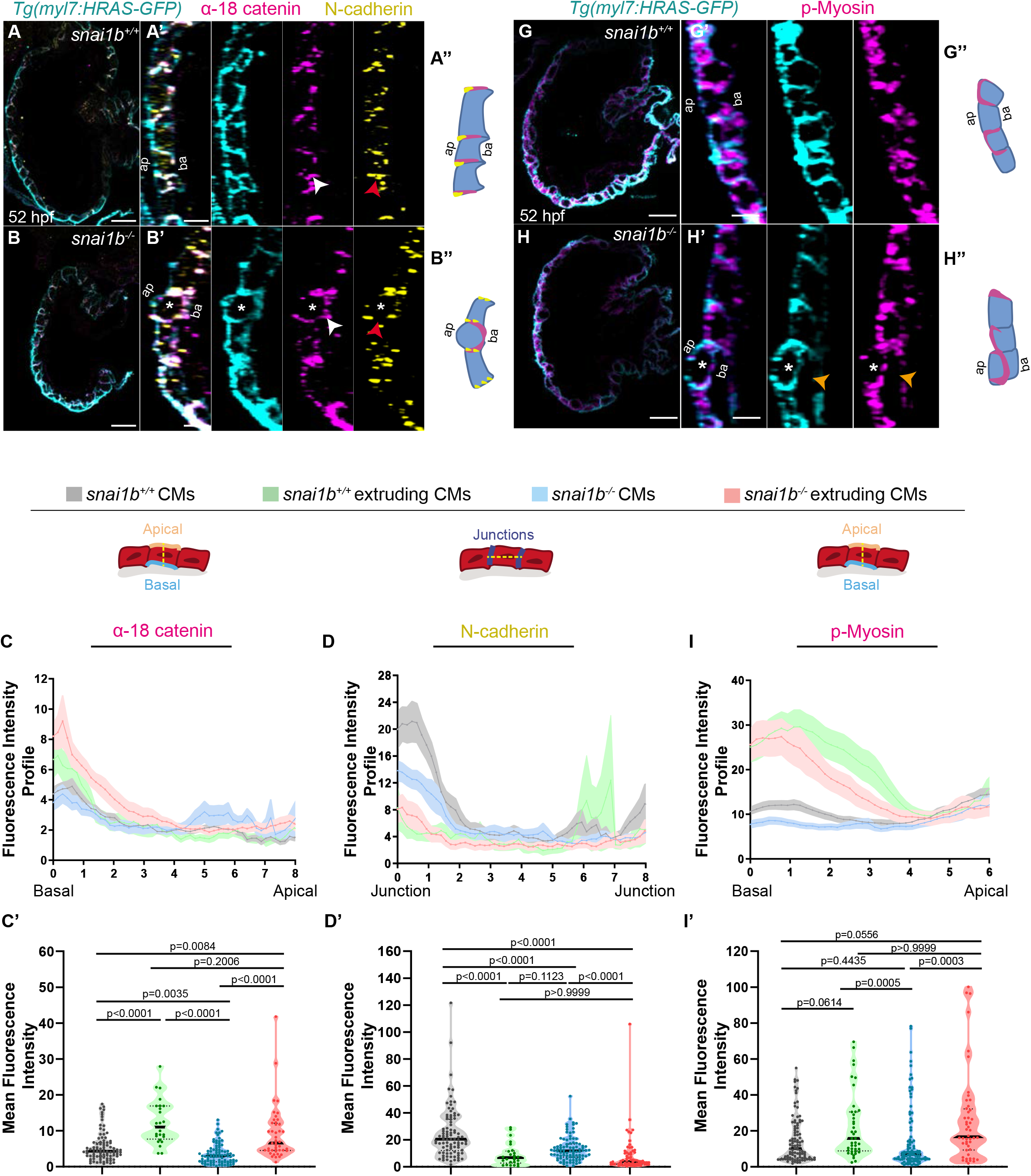
Extruding cardiomyocytes exhibit changes in actomyosin contractility. **A-B”)** Representative images of *snai1b*^+/+^ heart (A) immunostained for α-18 catenin, N-cadherin and GFP compared with mutant sibling (B) at 52 hpf. High magnification of CMs (A’,B’). Schematics illustrate the localization of opened α-catenin (magenta) in the basal domain in *snai1b*^-/-^ extruding CMs and of breaks of N-cadherin (yellow) in the junctional domain in *snai1b*^-/-^ CMs (A”-B”). **C-D’)** Fluorescence Intensity Profile (FIP) (C-D) and mean Fluorescence Intensity (mFI) (C’-D’) of α-18 catenin and N-cadherin immunostaining in *snai1b*^+/+^ and *snai1b*^-/-^ CMs, and in *snai1b*^+/+^ and *snai1b*^-/-^ extruding CMs. The α-18 catenin epitope is observed in the basal domain (white arrows in B’) of extruding CMs (white asterisks in B’) in *snai1b* mutants, and a reduction in junctional N-cadherin (red arrowheads in B’) is observed in *snai1b*^-/-^ mutant CMs. (FIP α-18 catenin: *snai1b* CMs, N=179; *snai1b*^+/+^ extruding CMs, N=60; *snai1b*^-/-^ CMs, N=140; *snai1b*^-/-^ extruding CMs, N=54; mFI α-18 catenin: *snai1b*^+/+^ CMs, N=90; *snai1b*^+/+^ extruding CMs, N=24; *snai1b*^-/-^ CMs, N=88; *snai1b*^-/-^ extruding CMs, N=44. FIP N-cadherin: *snai1b*^+/+^ CMs, N=90; *snai1b*^+/+^ extruding CMs, N=12; *snai1b*^-/-^ CMs, N=98; *snai1b*^-/-^ extruding CMs, N=49; mFI N-cadherin: *snai1b*^+/+^ CMs, N=90; *snai1b*^+/+^ extruding CMs, N=25; *snai1b*^-/-^ CMs, N=92; *snai1b*^-/-^ extruding CMs, N=70). **G-H**’’) Representative images of *snai1b* mutant heart (H) immunostained for p-Myosin and GFP compared with wild-type sibling heart (G) at 52 hpf. High magnification of CMs (G’,H’). Schematics illustrate the basal enrichment of p-Myosin (magenta) in extruding CMs in *snai1b* mutants (G”-H”). **I-J)** FIP (I) and mFI (I’) of p-Myosin immunostaining in *snai1b*^+/+^ and in *snai1b*^-/-^ CMs, and *snai1b*^+/+^ and *snai1b*^-/-^ extruding CMs. p-Myosin is enriched basally (orange arrows in H’) in extruding CMs in *snai1b* mutants (white asterisks in H’). (FIP p-Myosin: *snai1b*^+/+^ CMs, N=204; *snai1b*^+/+^ extruding CMs, N=60; *snai1b*^-/-^ CMs, N=140; *snai1b*^-/-^ extruding CMs, N=49; mFI p-Myosin: *snai1b*^+/+^ CMs, N=100; *snai1b*^+/+^ extruding CMs, N=29; *snai1b*^-/-^ CMs, N=153; *snai1b*^-/-^ extruding CMs, N=48). Plot values represent means ± S.E.M. (C,D,I). In the violin plots (C’,D’,I’), solid black lines indicate median. p-values determined by Kruskal-Wallis test (C’,D’,I’). Scale bars: 20 μm (A,B,G,H); 2 μm (A’,B’,G’,H’). Ap, apical; ba, basal; N, number of CMs.

### Intermediate filament gene expression is dysregulated in *snai1b*^-/-^ hearts

To further understand how the transcription factor Snai1b is required to maintain myocardial wall integrity, we compared the *snai1b*^+/+^ and *snai1b*^-/-^ cardiac transcriptomes at 48 hpf, a time when CM extrusion is starting to be observed (Figure 3A). Since Snai1 primarily acts as a transcriptional repressor^32^, we focused on the genes upregulated in *snai1b*^-/-^ hearts compared with wild type. Out of the 339 upregulated genes, gene ontology analysis revealed an enrichment of genes related to the cytoskeleton (Figure S4A), particularly an upregulation of intermediate filament (IF) genes (Figure 3B). In the myocardium, IF protein networks have been linked with diverse functions including mechanochemical signal transduction and regulation of adhesion and migration^33^. In particular, mutations that modify posttranslational modification sites in IF proteins have been associated with cardiomyopathy^34^, but how IF genes are regulated at the transcriptional level remains poorly understood. Interestingly, the muscle-specific IF gene *desmin b* was upregulated in *snai1b*^-/-^ hearts (Figure 3C), further indicating that Snai1 modulates CM development cell-autonomously. In muscle cells, Desmin is localized to Z-discs and desmosomes within intercalated discs, and an imbalance of Desmin protein is a major cause of cardiomyopathies^35^. Using quantitative PCR and immunostaining to analyze *desmin* at the RNA and protein levels, respectively, we first confirmed the upregulation of *desmin b* transcript and Desmin protein in *snai1b*^-/-^ hearts at 48 and 52 hpf (Figures 3D and 3G–3I’). Notably, extruding *snai1b*^-/-^ CMs exhibit an enrichment of Desmin in their basal domain and a correlative loss of Desmin at intercellular junctions (Figures 3I–3I’), indicating an abnormal localization of Desmin. As IFs are known to regulate actomyosin contractility in keratinocytes and astrocytes^36^, these data suggest that basal enrichment of Desmin promotes CM extrusion in *snai1b*^-/-^ hearts. To further test the hypothesis that Snai1 represses *desmin b* expression, we analyzed *desmin* transcript levels upon *snai1b* overexpression. qPCR analysis 4.5 hours after mRNA injection confirmed downregulation of *desmin b* transcript levels when *snai1b* was overexpressed (Figures S4B–S4C) compared with *gfp* injected controls. Similarly, qPCR analysis of hearts that overexpress CM-specific *snai1b* showed a reduction of *desmin b* transcript levels by 40% (Figures 3E–3F). Taken together, these data indicate that Snai1b regulates Desmin in two ways: 1) transcriptionally, whereby it represses *desmin b* expression, and 2) post-translationally, whereby it regulates Desmin localization at cellular junctions.

**Figure 3.**
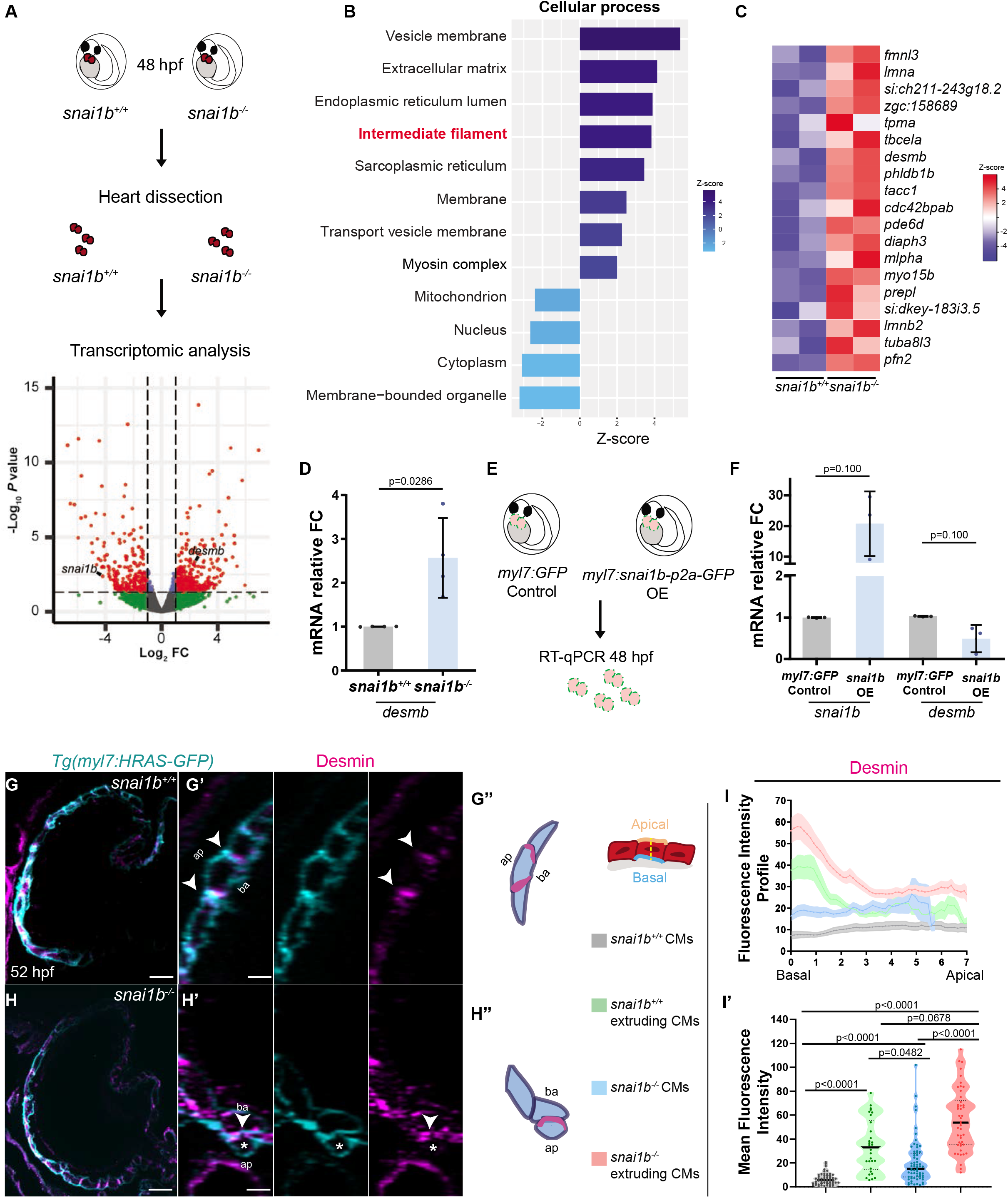
Transcriptomic analysis reveals upregulation of intermediate filament genes in *snai1b* mutant hearts. **A)** RNA extracted from 48 hpf *snai1b*^+/+^ and *snai1b*^-/-^ hearts was used for transcriptomic analysis. **B)** GO analysis of cellular processes shows enrichment of intermediate filament components. **C)** Heatmap of a list of upregulated cytoskeletal genes, including *desmin b*. **D)** Relative mRNA levels of *desmin b* are significantly increased in *snai1b*^-/-^ hearts at 48 hpf; n=4 biological replicates, 30 hearts each. **E)** Schematic of *snai1b* overexpression under a *myl7* promoter; *snai1b* and *desmin b* mRNA levels analyzed at 48 hpf. **F)** Relative mRNA levels of *desmin b* are significantly reduced in *snai1b* myocardial-specific overexpressing hearts at 48 hpf; n=3 biological replicates, 30 hearts each. **G-H**”) Representative images of 52 hpf *snai1b* mutant heart (H) immunostained for Desmin and GFP compared with wild type (G). High magnification of CMs (G’,H’). Schematics (Desmin in magenta) illustrate the basal localization of Desmin in *snai1b*^-/-^ extruding CMs (G”-H”). **I-I’)** FIP (I) and mFI (I’) of Desmin in *snai1b*^+/+^ and *snai1b*^-/-^ CMs, and *snai1b*^-/-^ extruding CMs. Desmin immunostaining is observed throughout the *snai1b*^-/-^ CM, with an enrichment in the basal domain (white arrowheads in H’-G’) in extruding CMs (white asterisks in G’). (FIP: *snai1b*^+/+^ CMs, N=49; *snai1b*^+/+^ extruding CMs, N=41; *snai1b*^-/-^ CMs, N=45; *snai1b*^-/-^ extruding CMs, N=41; mFI: *snai1b*^+/+^ CMs, N=56; *snai1b*^+/+^ extruding CMs, N=30; *snai1b*^-/-^ CMs, N=65; *snai1b*^-/-^ extruding CMs, N=46). Plot values represent means ± S.D. (D,F) or mean ± S.E.M. (I). In the violin plot (I’), solid black lines indicate median. p-values determined by Mann-Whitney *U* (D,F) or Kruskal-Wallis test (I’). Scale bars: 20 μm (G,H); 2 μm (G’,H’). Ap, apical; ba, basal; n, number of embryos; N, number of CMs; FC, fold change.

### *desmin b* overexpression in CMs promotes their extrusion

Both loss^37,38^ and gain^39^ of Desmin expression have been associated with cardiac defects. Thus, we asked whether an imbalance in *desmin b* expression could lead to CM extrusion by overexpressing *desmin b* mosaically in CMs. We observed that *desmin b* overexpressing CMs were more prone to extrude compared with *gfp* overexpressing CMs (Figures 4A–4C), suggesting that IFs are needed at their endogenous levels to maintain myocardial wall integrity. We hypothesized that increased Desmin levels induce CM extrusion by disrupting desmosome organization leading to compromised cell-cell adhesion and/or by increasing cell contractility basally. To test these hypotheses, we first used electron microscopy to analyze desmosomes at the ultrastructural level, but observed no obvious defects in *snai1b*^-/-^ CMs compared with wild type (Figures S4F–S4I). This result is consistent with a previous study that shows intact desmosomes in extruding epithelial cells^40^. Second, to test whether accumulation of Desmin in the basal side of CMs was associated with increased cell contractility, we immunostained *desmin b* overexpressing CMs with the α-catenin α-18 antibody. Notably, *desmin b* overexpressing CMs exhibited an accumulation of the α-catenin α-18 epitope basally (Figures 4D–4E’), suggesting that increased Desmin correlates with increased actomyosin-based cellular contractility basally and apical extrusion.

**Figure 4.**
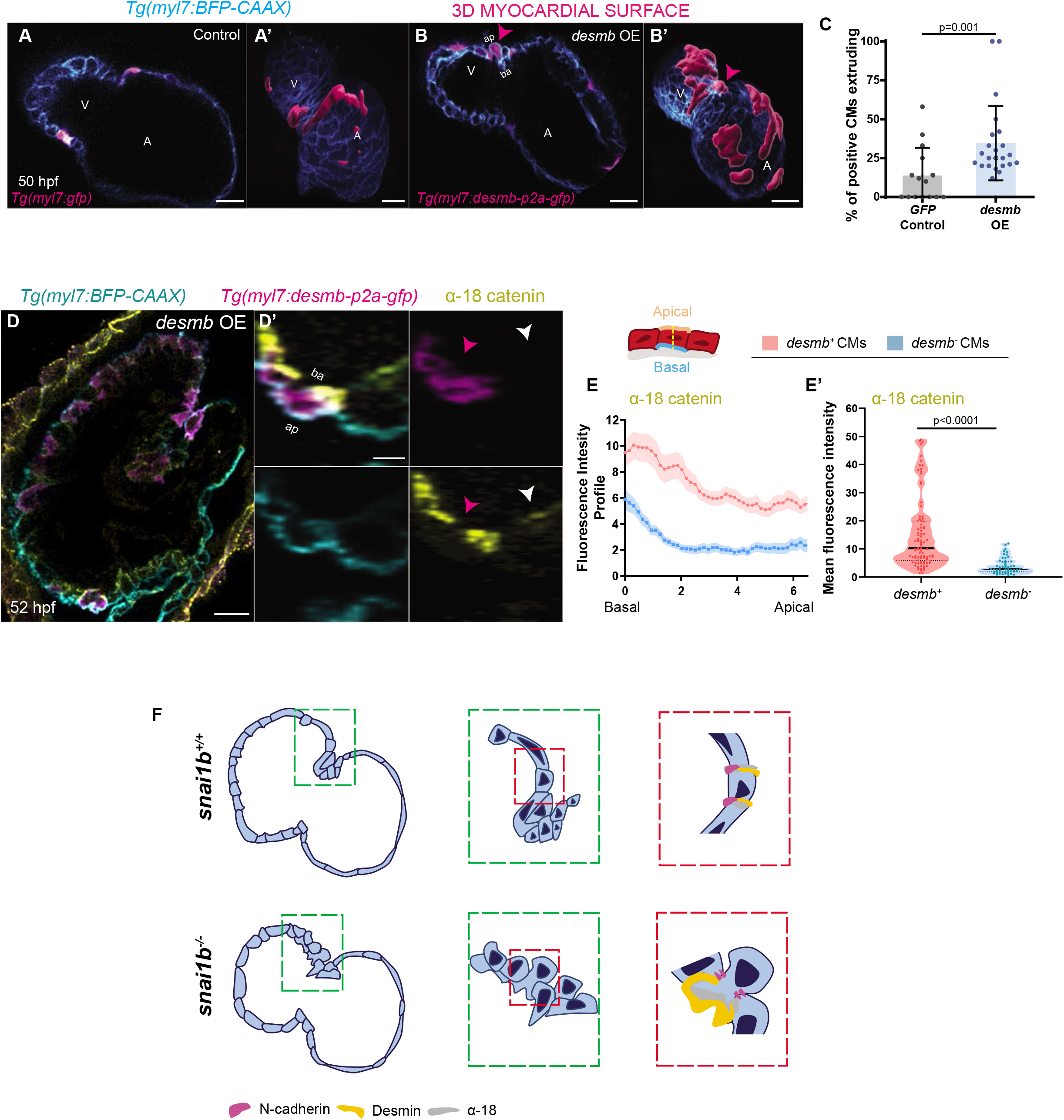
*desmin b* overexpression in cardiomyocytes induces their extrusion. **A-B**’) Single-plane images of wild-type embryos injected with *myl7:GFP* (A, 3D surface rendering of GFP+ CMs in A’) or with *myl7:desmb-p2a-GFP* (B, 3D surface rendering of *desmin b+* CMs in B’) at 50 hpf. **C)** A higher percentage of CMs extrude when overexpressing *desmin b* (n=23) compared with control (n=15) (magenta arrowheads in B, B’). **D-D**’) Representative image of the heart of embryos injected with *myl7:desmb-p2a-GFP* (D) immunostained for α-18 catenin, GFP and BFP at 52 hpf. High magnification of CMs (D’). **EE’)** FIP (E) and mFI (E’) of α-18 catenin in CMs that overexpress *desmin b* (magenta arrowhead) and CMs that do not overexpress *desmin b* (white arrowhead) CMs. α-18 catenin epitope immunostaining is enriched in the basal domain in *desmin b* overexpressing CMs. (FIP: *desmin b+* CMs, N=179; *desmin b-* CMs, N=60; mFI: *desmin b+* CMs, N=59; *desmin b-* CMs, N=56). **F)** Model: loss of *snai1b* leads to CM extrusion and basal enrichment of the actomyosin machinery and Desmin, thereby affecting myocardial wall integrity. Plot values represent means ± S.D. (C) or means ± S.E.M. (E). In the violin plot (E’), solid black lines indicate median. p-values determined by Mann-Whitney *U* test. Scale bars: 20 μm (A-B’,D); 2 μm (D’). V, ventricle; A, atrium; ap, apical; ba, basal; n, number of embryos.

## Conclusions

A role for Snai1 in cell extrusion has been reported in *Drosophila* embryos as well as in mammalian cell culture. During gastrulation in *Drosophila*, Snai1 promotes the medio-apical pulsations of contractile Myo-II that drive apical constrictions^9, 43, 44^. However, the identity of the transcriptional targets of Snai1 that promote cellular contractility in this system remains unknown. Recent *in vitro* studies have reported Snai1-mediated upregulation of active RhoA at the mRNA and protein levels, leading to increased cortical actomyosin and apical extrusion^11^. Here, our work uncovers a previously unsuspected role for the EMT-inducing factor Snai1 in limiting CM extrusions by regulating IF gene expression. We show that the CM extrusions in *snai1b* mutants are associated with increased accumulation of actomyosin basally, providing more evidence for a role of Snai1 in regulating cell contractility through actin networks. This function appears to be partly independent of Snai1’s role in EMT, as no obvious changes in localization of the apical marker Podocalyxin were observed. Our data clearly show the requirement of Snai1 in maintaining epithelial tissue integrity in a vertebrate organ, and add to the growing evidence that Snai1 has EMT-independent roles in epithelial tissues.

Furthermore, we report a previously uncharacterized function of Snai1 in regulating *desmin* expression and find that an increase in Desmin levels perturb tissue cohesiveness. Although IFs including Vimentin^43^ and Keratin^42, 46^ are known to accumulate at the interface between extruding cells and their neighbors, our study provides evidence that increased Desmin levels in CMs are correlated with mislocalization of the actomyosin machinery in the basal domain and cell extrusion. These data are consistent with previous findings that IFs can regulate the localization and activation of the actomyosin network, with factors such as Vimentin directly binding to actin and modulating RhoA activity^45^, and Keratin directly binding to Myosin^46^. In addition to a role for Desmin in maintaining nuclear membrane architecture in CMs^47^, our results shed light on the function of Desmin in preventing CM extrusion and maintaining myocardial integrity.

Our results also uncover the requirement of Snai1 and correct levels of Desmin in maintaining myocardial integrity under contraction-induced mechanical pressure. Heart contraction is essential in patterning the cardiac tissue: without heartbeat, cardiac valves and trabecular networks fail to form^50, 51^. However, our results indicate that without a strong intracellular cytoskeletal network regulated by Snai1, the heartbeat-induced mechanical forces can lead to an increase in cell extrusion. While it has been shown that an increase in cell contractility due to changes in morphology, adhesion or cell density drives cell extrusion^23, 24, 26, 28, 29, 52, 53^, we present evidence that external mechanical forces contribute to non-apoptotic CM extrusion during cardiac development, and that actomyosin and IF cytoskeletal regulation is critical to prevent CM extrusion.

In conclusion, our findings uncover molecular mechanisms that suppress cell extrusion in a tissue under constant mechanical pressure, and show a multifaceted, context-dependent role for Snai1 in promoting EMT^2^, and also in maintaining tissue integrity during vertebrate organ development (Figure 4F).

## Acknowledgement

This work was supported by funds from the Max Planck Society to D.Y.R.S.; a European Molecular Biology Organization (EMBO) Advanced Fellowship (ALTF 642-2018) and a Canadian Institute for Health Research Fellowship (293898) to F.G.; and an EMBO fellowship (LTF 1569-2016), a Humboldt fellowship and a Cardio-Pulmonary Institute Grant (EXC 2026, project ID 390649896) to R.P. We would like to thank Michelle Collins, Paolo Panza, Chi-Chung Wu, Mridula Balakrishnan, Srinivas Allanki, Giulia Boezio and Simon Perathoner for comments on the manuscript, and Prof. Akira Nagafuchi for the α-18 antibody.

## Author contributions

Conceptualization, A.G., F.G., R.P., A.B.B., and D.Y.R.S.; Methodology, A.G, F.G., J.P., S.G.; Validation, A.G.; Formal Analysis, A.G., S.G.; Investigation, A.G., J.P., S.G.; Writing – Original Draft, A.G., F.G. and D.Y.R.S.; Writing – Reviewing & Editing, all; Visualization, A.G.; Supervision, F.G., A.B.B. and D.Y.R.S; Project Administration, D.Y.R.S.; Funding Acquisition, D.Y.R.S.

---

**Figure S1.**
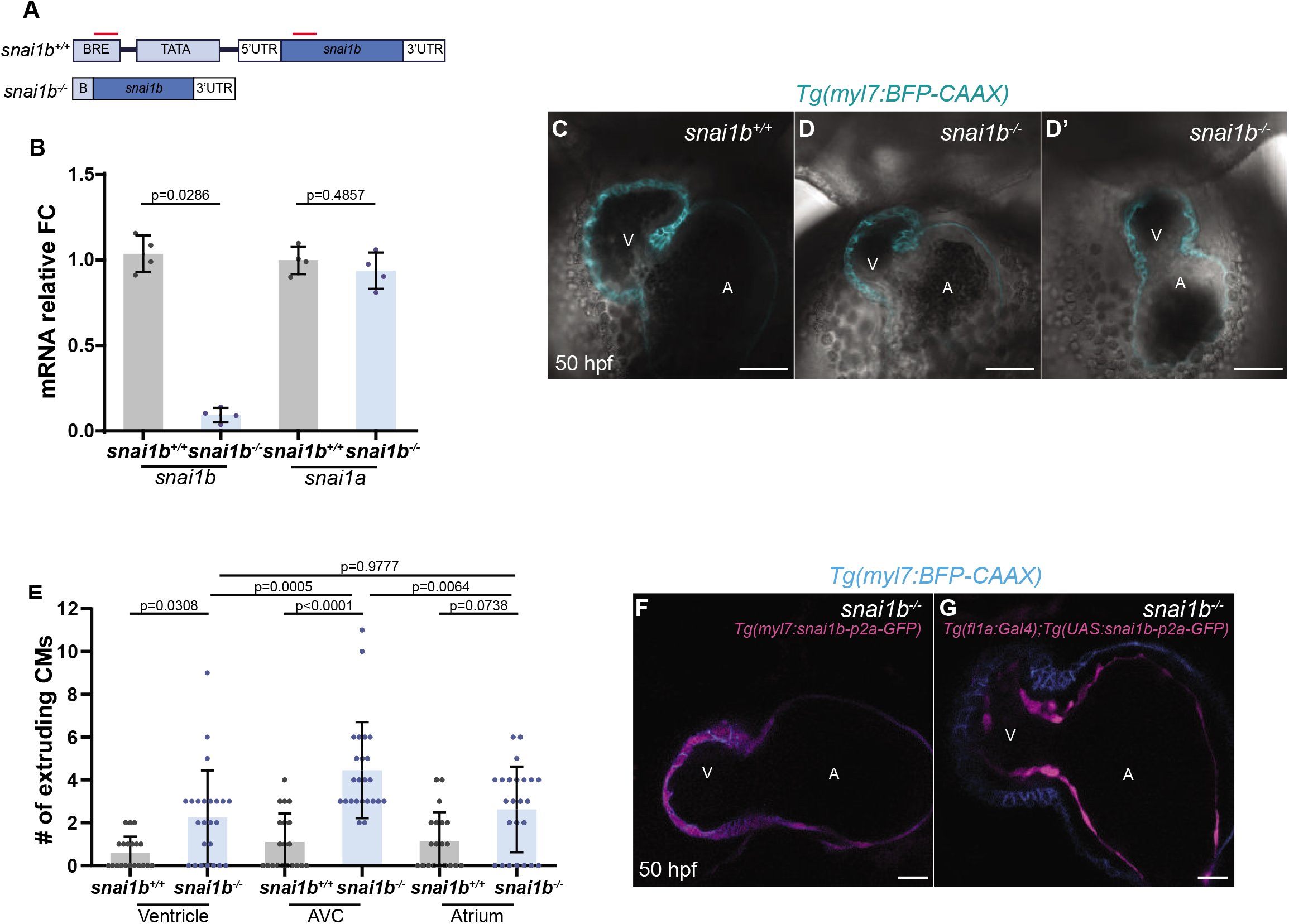
Expression pattern of *snai1b* during zebrafish cardiac development, and generation of *snai1b* mutants. **A)** Schematic of the promoter-less *snai1b* allele. Two gRNAs (red lines) were used to generate *snai1b* mutant allele lacking 1300 bp upstream of the start codon. **B)** Relative mRNA levels of *snai1b* are significantly reduced in *snai1b*^-/-^ *hearts* at 48 hpf, whereas *snai1a* expression levels are unchanged, indicating lack of transcriptional adaptation by the paralogue; n=4 biological replicates, 30 embryos each. **C-D’)** Single-plane images of *snai1b*^+/+^ (C) and *snai1b*^-/-^ (D-D’) hearts at 50 hpf, with 50% of *snai1b*^-/-^ mutant hearts exhibiting cardiac looping defects. **E)** The majority of extruding *snai1b*^-/-^ CMs are localized in the AVC compared with the ventricle and the atrium (*snai1b*^+/+^, n =20; *snai1b*^-/-^, n=24). **F-G)** Single–plane images of *snai1b*^-/-^ hearts overexpressing *snai1b* under a *myl7* promoter (F) or a *fli1a* promoter (G). Plot values represent means ± S.D.; p-values determined by Mann-Whitney *U* test (B) or by one-way ANOVA followed by multiple comparisons with Dunn test (E). Scale bars: 20 μm (C-D’,F,G). BRE, B recognition element; V, ventricle; A, atrium; n number of embryos.

**Figure S2.**
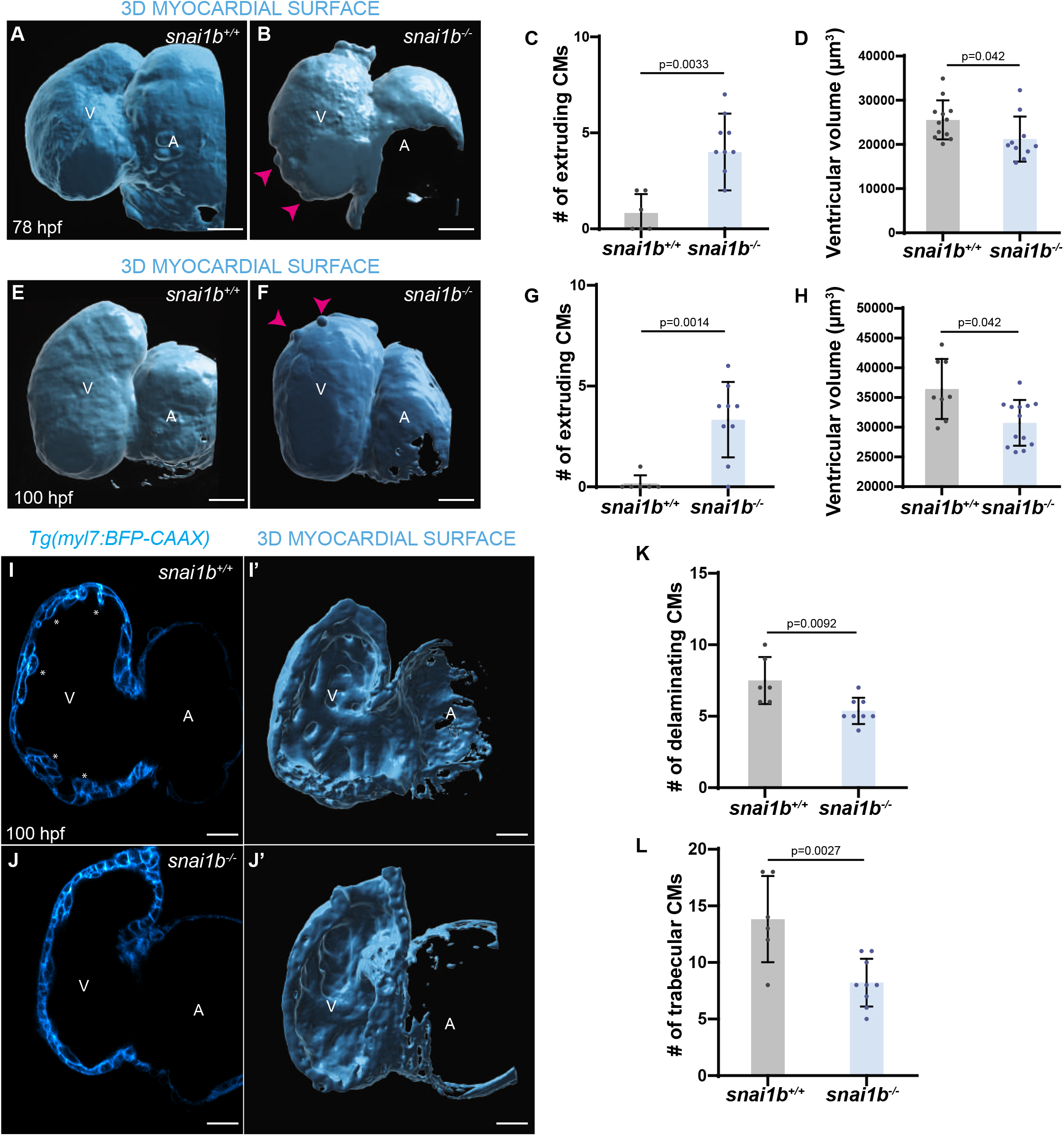
*snai1b* mutants exhibit reduced cardiac trabeculation and heart rate. **A-H)** 3D surface rendering of the ventricle at 78 and 100 hpf. *snai1b*^-/-^ larvae (B, F) exhibit extruding CMs (magenta arrowheads) and reduced ventricular volume compared with the wild-type larvae (A, E). Increased CM extrusion numbers (C, G) and significantly reduced ventricular volumes (D, H) are observed in *snai1b*^-/-^ larvae compared with wild types at 78 and 100 hpf (C, *snai1b*^+/+^, n=6; *snai1b*^-/-^, n=10; G, *snai1b*^+/+^, n=12; *snai1b*^-/-^, n=10; D, *snai1b*^+/+^, n=6; *snai1b*^-/-^, n=9; H, *snai1b*^+/+^, n=8; *snai1b*^-/-^ n=13). **I-J’)** Single-plane images and 3D surface rendering of trabecular CMs (asterisks in I) of *snai1*^-/-^ larvae (J-J’) and wild type (I-I’) at 100 hpf. **K-L)** Decreased delaminating (K) and trabecular (L) CM numbers are observed in *snai1b*^-/-^ larvae compared with wild type at 78 (*snai1b*^+/+^, n=6; *snai1b*^-/-^, n=8) and 100 hpf (*snai1b*^+/+^, n=6; *snai1b*^-/-^, n=9). Plot values represent means ± S.D.; p-values determined by Student’s t-test (C,D,G,H,K,L). Scale bars: 20 μm. V, ventricle; A, atrium; n, number of embryos.

**Figure S3.**
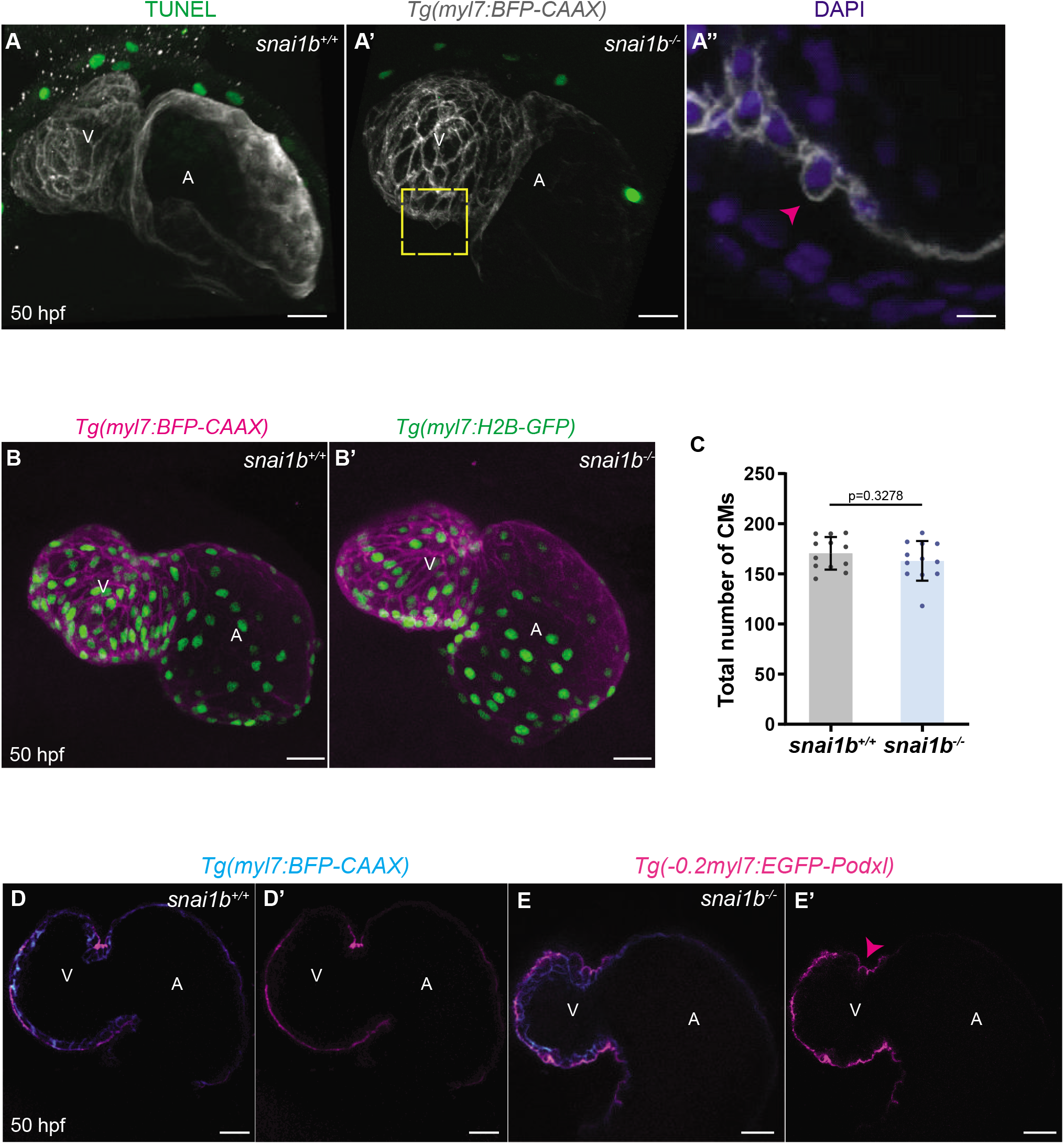
*snai1b* mutants exhibit wild-type like cardiomyocyte numbers. **A-A”’)** TUNEL assays indicates that extruding CMs in *snai1b*^-/-^ are not apoptotic. Maximum intensity projection of *snai1b*^+/+^ (A) and *snai1b*^-/-^ hearts (A’-A”). Close-up of extruding CM (magenta arrowheads) labeled with DAPI, but not with TUNEL (A”). **B-B’)** Maximum intensity projections of *Tg*(*myl7:H2B-GFP*) hearts in wild types (B) and *snai1b*^-/-^ (B’) at 52 hpf. **C)** The total number of CMs does not change in *snai1b*^-/-^ (n=12) hearts compared with wild types (n=12). **D-E’)** Single-plane images of *Tg*(*−0.2myl7:EGFP-podocalyxin*) *snai1b*^+/+^ (D-D’) and *snai1b*^-/-^ (E-E’) hearts at 50 hpf. No changes in the localization of the apical marker Podocalyxin are observed in extruding CMs (magenta arrow in E’). Plot values represent means ± S.D.; p-value determined by Student’s t-test. Scale bars: 20 μm (A-A’,B-B’,D-E’); 10 μm (A”). V, ventricle; A, atrium; n, number of embryos.

**Figure S4.**
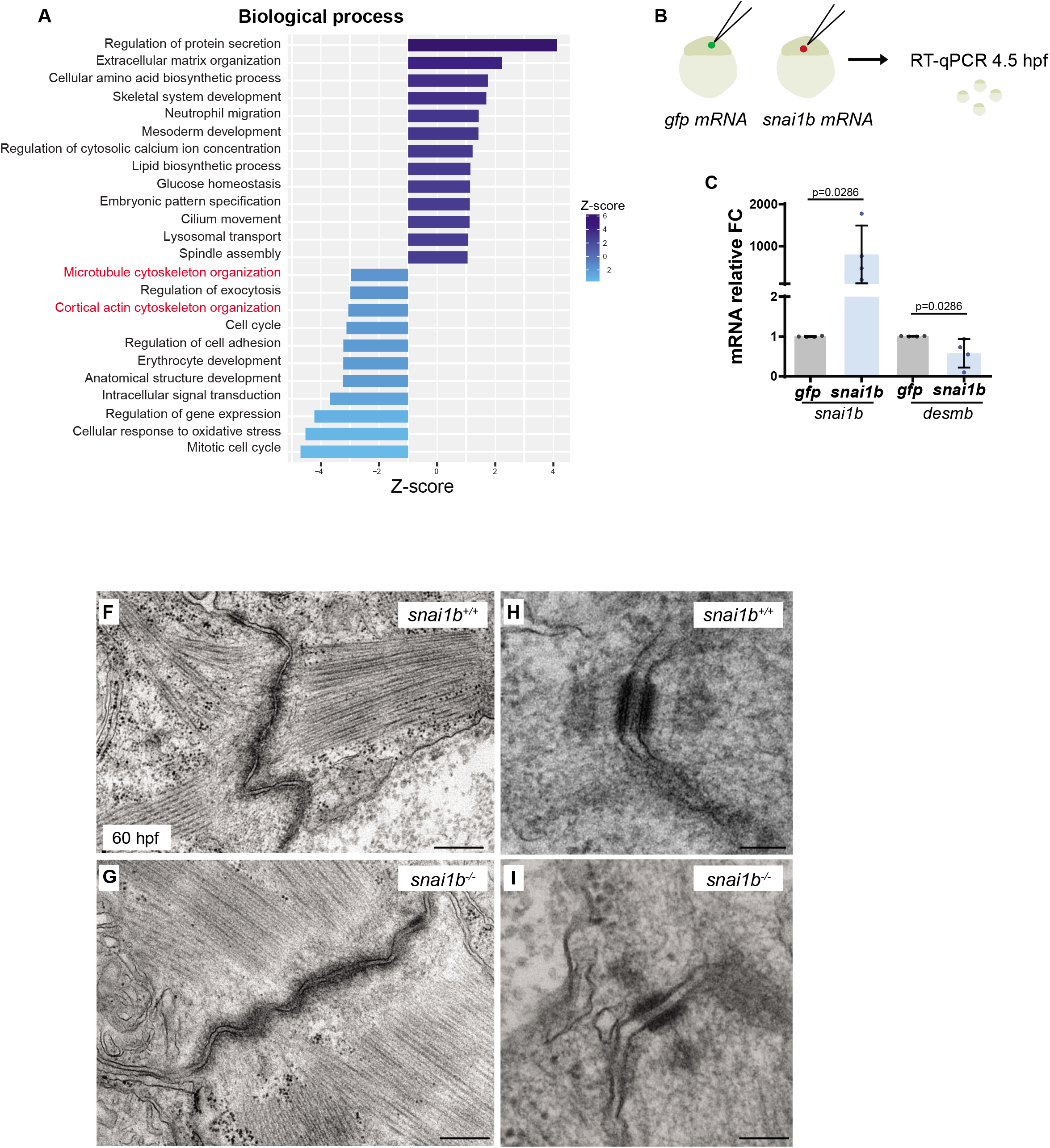
*snai1b* mRNA injections decrease *desmin b* mRNA levels. **A)** GO analysis of biological processes shows enrichment of cytoskeletal related genes (in red). **B-C)** Schematic of *gfp* and *snai1b* mRNA injections. *snai1b* and *desmin b* mRNA levels were analyzed in injected embryos at 4.5 hpf. Relative mRNA level of *snai1b* is significantly increased, whereas *desmin b* expression level is decreased at 4.5 hpf. n=4 biological replicates, 30 embryos each. **F-I)** Transmission electron microscopy (TEM) images showing the ultrastructure of fascia adherens (F, G) and desmosomes (H, I) in *snai1b*^+/+^ and *snai1b*^-/-^. No noticeable changes were observed in these structures. Plot values represent means ± S.D.; p-values determined by Mann-Whitney *U*. Scale bars: 200 nm (F,G); 500 nm (H,I).

